# Unraveling Wheat Aphid Diversity in Bangladesh Through Integrative Genetic and Morphological Approaches

**DOI:** 10.64898/2026.07.27.741099

**Authors:** Md. Moshiur Rahman, Most Sirajum Munira, Md Forhad Hossen, Md Nizam Uddin, Mostafizur R. Shah

**Author notes:** Corresponding author (MMRS). These authors contributed equally to this work and share first authorship (Md. Moshiur Rahman, Most Sirajum Munira).

## Abstract

Accurate species identification is fundamental for understanding insect pest diversity and developing effective management strategies. Wheat aphids are among the most destructive insect pests worldwide, yet comprehensive molecular confirmation of species diversity in Bangladesh has been lacking. This study employed an integrative taxonomic approach combining DNA barcoding and morphological characterization to identify wheat aphid species collected from major wheat-growing regions of Bangladesh. Aphid specimens were sampled from 11 locations across four districts during the 2024-2025 wheat growing season. A partial fragment of the mitochondrial cytochrome *C* oxidase subunit I (COI) gene was amplified and sequenced, followed by BLAST analysis, phylogenetic reconstruction using the Maximum Likelihood method, genetic distance estimation, and detailed morphometric analyses. COI sequence analysis identified five wheat aphid species: *Sitobion avenae*, *Rhopalosiphum padi*, *R. maidis*, *R. rufiabdominalis*, and *Hysteroneura setariae*. *Sitobion avenae* was the predominant species, occurring across all surveyed regions. BLAST similarity ranged from 99.68% to 100%, confirming reliable species-level identification. Phylogenetic analysis grouped all specimens into five well-supported species-specific clades with strong bootstrap support (97-100%), corroborating molecular identification. Pairwise genetic distance analysis revealed very low intraspecific divergence (0.0–0.18%) and clear interspecific divergence (4.9-11.9%), indicating strong genetic discrimination among species. Morphological and morphometric analyses further distinguished the five species, with significant interspecific differences (*P* < 0.0001) observed for body length, antennal length, tubercle width, cauda length, and cornicle length. The concordance between molecular and morphological evidence demonstrates the robustness of the integrative taxonomic framework. This study provides the first comprehensive molecular and morphological characterization of wheat aphid species in Bangladesh, establishing a reliable baseline for biodiversity assessment, pest surveillance, insecticide resistance monitoring, and the development of evidence-based integrated pest management strategies to support sustainable wheat production amid changing agroecological conditions.

## INTRODUCTION

Wheat (*Triticum aestivum* L.) supplies approximately 20% of global human caloric intake, making it the second most important staple food crop after rice in Bangladesh [1]. In Bangladesh, wheat occupies ∼0.3 million hectares during the winter (*rabi*) season (November–March). It serves as a vital nutritional and economic addition to rice-based farming systems [2,3]. Among the threats to wheat production, aphids (Hemiptera: Aphididae) are the most significant pests affecting cereal crops worldwide [4]. These sap-sucking insects harm wheat by extracting nutrients, which deplete photosynthates and reduces grain filling. They also promote the growth of sooty mold through honeydew secretion and spread harmful plant viruses like Barley Yellow Dwarf Virus (BYDV). This virus can lower wheat yield by up to 80% in susceptible systems [4,5,6]. In South Asia, major aphid outbreaks during tillering to grain filling routinely depress wheat yields by 15–40% [7,8]. Climate change is exacerbating this threat, as rising winter temperatures and altered cropping patterns favor prolonged aphid reproduction and dispersal [4]. These trends create an urgent need to understand aphid species diversity and distribution in Bangladesh’s major wheat-growing zones.

Accurate species identification is the cornerstone of ecologically informed pest management. It supports monitoring programs, resistance checks, biological control methods, and targeted integrated pest management (IPM) strategies. Traditional aphid classification relies on physical traits and host plant relationships [9,10]. However, this approach is limited by the significant physical variation seen in the Aphididae family. Differences in phenotype caused by host adaptation, geographical separation, reproduction methods, and varied life cycles often create overlapping traits among similar species, resulting in unstable classifications and hidden species complexes [11,12]. This is especially evident in important cereal-related genera, where several species, such as *Rhopalosiphum maidis*, *R. padi*, *Sitobion avenae*, and *Schizaphis graminum*, share similar physical traits that complicate reliable identification [10,13]. Reliance solely on morphological keys therefore risks inaccurate species delimitation and compromises the integrity of biodiversity records [14,15].

Integrative taxonomy, which combines molecular and morphological data, has become the standard approach to tackle the challenges in aphid classification. The mitochondrial cytochrome *C* oxidase subunit I (COI) gene is recognized as the universal DNA barcode for insect identification due to its strong ability to differentiate species across various groups [16,17,18]. In aphids, using COI for barcoding has proven to be very effective in uncovering hidden diversity. Studies have shown a significant mismatch between morphological classifications and molecular lineages in various subfamilies. Recent research on Korean aphids confirmed that COI barcoding can reliably distinguish most morphospecies while also revealing cryptic genetic lineages that are not detectable by physical traits alone [28]. Regional molecular surveys for cereal aphids have progressed in India, Pakistan, and China [8,19,20]. These studies have improved the understanding of the relationships among *Sitobion*, *Rhopalosiphum*, and *Schizaphis* populations across Asia. Together, they demonstrate the importance of integrating both morphological and molecular data for a thorough assessment of aphid diversity and pest monitoring.

Despite the recognized agricultural significance of wheat in Bangladesh and the regular aphid problems in its main growing areas, the diversity of wheat aphids in the country has not been studied using molecular techniques. Existing records are based exclusively on morphological observation, leaving species identities genetically unvalidated and potentially incomplete. There are no reference barcode sequences for Bangladeshi wheat aphids in major global databases like GenBank and BOLD Systems, which is a major gap in the understanding of aphids in South Asia. Bangladesh has several unique agroecological zones for wheat cultivation, which vary widely in temperature, humidity, cropping practices, and landscape characteristics. These environmental factors are known to affect aphid community structure and genetic makeup [8,13]. Yet no comparative assessment of aphid species diversity or population genetics across these zones has been conducted, constituting both a geographic and a taxonomic gap in regional cereal pest research. This study aims to fill these gaps using an integrative approach that combines detailed physical characterization with COI-based DNA barcoding. We hypothesize that multiple species of the genera *Sitobion*, *Rhopalosiphum*, and *Schizaphis* infest wheat fields in Bangladesh, and that molecular analysis will reveal hidden genetic diversity and geographical patterns that cannot be detected through physical traits alone. The specific objectives were to: (i) accurately identify wheat aphid species using combined morphological and molecular evidence; (ii) characterize genetic variability and infer phylogenetic relationships among collected populations relative to Asian reference sequences; and (iii) establish the first genetically validated barcode reference database for wheat-associated aphids in Bangladesh. This research represents the first integrated taxonomic study of wheat aphids in Bangladesh and will provide data for molecular diagnostics, pest monitoring, and informed aphid management in South Asian cereal farming systems.

## MATERIALS AND METHODS

### Study area and sampling

Field surveys were conducted during the 2024–2025 rabi season (October 2024 to April 2025) across four foremost wheat-growing districts of Bangladesh, namely Dinajpur, Thakurgaon, Panchagarh, and Rajshahi. These districts represent the principal agroecological wheat production zones of the country and differ in climatic conditions, including temperature, relative humidity, and cropping intensity, thereby providing broad geographic coverage for assessing aphid diversity.

Within each district, 2–4 wheat fields separated by at least 2 km were surveyed. In each field, 10–15 wheat plants were randomly selected along a diagonal transect for aphid collection. A total of 11 geographically distinct sampling localities were included in the study, and each locality was considered an independent population for molecular and morphometric analyses. All laboratory experiments and analyses were conducted at the Insect Genomics Laboratory of the Bangladesh Wheat and Maize Research Institute.

### Aphid collection and preservation

Aphid specimens comprising both adults and nymphs were collected from wheat plants at different growth stages ranging from tillering to grain filling. Specimens were systematically collected from the adaxial and abaxial leaf surfaces, stems, and spikes using a fine camel-hair brush and immediately transferred into sterile pre-labeled 1.5 mL microcentrifuge tubes. Geographic coordinates of each sampling site were recorded using a Garmin eTrex 10 handheld GPS receiver. Detailed information regarding specimen codes, host plant, collection locality, and geographic coordinates is provided in Table 1.

**Table 1.** A comprehensive list of Aphid samples was generated from four major wheat growing districts of Bangladesh

| Sample No. | Sample ID | Location | Host | GPS |
| --- | --- | --- | --- | --- |
| 1 | WA-DnJ-A | Dinajpur | Wheat | 25°38'11.6664'' |
| 2 | WA-DnJ-B | Dinajpur | Wheat | 25°38'11.6664'' |
| 3 | WA-DnJ-C | Dinajpur | Wheat | 25°38'11.6664'' |
| 4 | WA-DnJ-D | Dinajpur | Wheat | 25°38'11.6664'' |
| 5 | WA-DnJ-E | Dinajpur | Wheat | 25°38'11.6664'' |
| 6 | WA-Tha-A | Thakurgaon | Wheat | 26°28'11.60'' |
| 7 | WA-Tha-B | Thakurgaon | Wheat | 26°28'11.60'' |
| 8 | WA-Raj-A | Rajshahi | Wheat | 24°22'23.911'' |
| 9 | WA-Raj-B | Rajshahi | Wheat | 24°22'23.911'' |
| 10 | WA-Pan-A | Panchagarh | Wheat | 26°20'7.3572'' |
| 11 | WA-Pan-B | Panchagarh | Wheat | 26°20'7.3572'' |

Specimens were divided into two preservation groups according to downstream analyses. Samples designated for morphological examination were preserved in 70% ethanol (v/v), whereas specimens intended for molecular analyses were immediately preserved in absolute ethanol (100%) to minimize DNA degradation. All samples were stored at −20°C until processing.

### Extraction of Genomic DNA

Total genomic DNA was extracted from five adult apterous females per population using the Wizard® Genomic DNA Purification Kit (Promega, Madison, WI, USA) following the manufacturer’s protocol with slight modification. Individual aphids were homogenized in 600 μl of Nuclei Lysis Solution using sterile plastic micropestles in 1.5 mL microcentrifuge tube and incubated at 65°C for 30 minutes. Following cell lysis, RNase treatment was performed according to the manufacturer’s instructions to remove residual RNA contamination. Proteins were precipitated using Protein Precipitation Solution, and genomic DNA was subsequently precipitated with room temperature isopropanol. DNA pellets were washed with 70% ethanol, air-dried at room temperature, and rehydrated in 100 μL DNA Rehydration Solution by overnight incubation at 4°C. DNA quantity and purity were assessed using a UV/Vis Nano Spectrophotometer (microDigital company Ltd). Samples with A₂₆₀/A₂₈₀ ratios ranging from 1.8 to 2.0 were considered suitable for downstream analyses. DNA integrity was further verified on 1% agarose gel stained with Ethidium Bromide. All DNA samples were normalized to 50 ng/μL prior to PCR amplification.

### PCR amplification of COI gene

A partial fragment of the mitochondrial cytochrome c oxidase subunit I (COI) gene was amplified using the universal primer pair LCO1490 (5′-GGTCAACAAATCATAAAGATATTGG-3′) and HCO2198 (5′TAAACTTCAGGGTGACCAAAAAATCA-3′) described by Folmer et al. [21]. PCR amplification was performed in a total reaction volume of 25 μL containing 12.5 μL DreamTaq Green PCR Master Mix (2×; Thermo Fisher Scientific), 1 μL of each primer (10 pmol/μL), 1 μL template DNA (50 ng/μL), and 9.5 μL nuclease-free water. Amplifications were conducted using MiniAmp Thermal Cycler (Applied Biosystems by ThermoFisher Scientific).

Thermal cycling conditions consisted of an initial denaturation at 95 °C for 2 minutes, followed by 40 cycles of denaturation of 95 °C for 30 seconds, annealing at 56.2 °C for 60 seconds, and extension at 72 °C for 30 seconds, with a final extension at 72 °C for 10 minutes. Each PCR run included a no-template negative control containing nuclease-free water and a positive control consisting of previously verified genomic DNA of *Sitobion avenae*. No amplification was detected in any negative control. PCR products were resolved on 0.8% agarose gel stained with Ethidium Bromide and visualized using Vilber Bio Print CX4 Gel Documentation System. A single band of ∼700 bp confirmed successful amplification.

### DNA sequencing and sequence processing

The crude PCR products were sent for sequencing at commercial facilities (OMC Ltd., Bangladesh) where the amplicons were sequenced in an automated sequencer (3500 Genetic Analyzer, Applied Biosystems, USA). Chromatograms were manually inspected, trimmed, and assembled into consensus sequences using Geneious Prime software (v2024.0.5; Biomatters Ltd., Auckland, New Zealand). Sequences were compared against the NCBI GenBank database using BLASTn. Sequences with ≥97% similarity were considered indicative of species-level identity [22].

### Multiple sequence alignment and phylogenetic analysis

COI sequences generated in this study, along with reference sequences retrieved from NCBI, were aligned using ClustalW implemented in MEGA 12 [23]. Ambiguous regions and gaps were removed, resulting in a final alignment of 572 bp. The best-fit nucleotide substitution model was selected using Bayesian Information Criterion (BIC) in MEGA 12. The Tamura 3-parameter model with invariant sites (T92+I) was identified as the best model [24]. Phylogenetic analysis was conducted using the Maximum Likelihood method with 1,000 bootstrap replications, and values ≥70% were considered strong support. *Aphis gossypii* was used as the outgroup taxon to root the phylogenetic tree due to its phylogenetic distinctness among cereal aphids. Pairwise genetic distances were estimated using the Kimura 2-parameter (K2P) model [25].

### Specimen preparation and slide mounting

Morphological voucher specimens were prepared for permanent slide mounting following the protocol of Noyes [26] with minor modifications. Prior to mounting, head, antennae, and wings were carefully detached and arranged on slides for proper visualization of diagnostic characters. Specimens were cleared in 10% KOH (6–12 h), neutralized in 10% glacial acetic acid for 10 min, and rinsed in distilled water for 10 min. Dehydration was performed using graded ethanol series (80%, 90%, 96%, and 100%). Specimens were cleared in clove oil and mounted in Canada balsam. Slides were dried at room temperature for 48 h. Voucher specimens were deposited in the Insect Genomics Laboratory Museum Collection, BWMRI (Voucher IDs: BWMRI-APH-01 to BWMRI-APH-11).

### Morphological Examination and Morphometric Analysis

Adult apterous females were examined for morphological and morphometric analyses. Specimens were observed using an Olympus SZX10 stereomicroscope equipped with an OLYMPUS DF PL 2X lens (Olympus Corporation, Tokyo, Japan). Species identification was performed using the diagnostic taxonomic keys of Blackman and Eastop [10]. Morphological identification was cross-validated with COI sequence data for integrative species confirmation. Morphometric traits including body length, antennal segments, cornicle length, and cauda length were measured using Olympus cellSens 4.3.1 version. Measurements were recorded in micrometers (μm) applying 1.0 magnification level.

### Statistical Analysis

All statistical analyses were performed in JMP Pro software version 13.0 [27]. Morphometric data were expressed as mean ± standard error (SE). Data meeting assumptions were analyzed without transformation. Differences among populations were assessed using one-way ANOVA followed by Tukey’s HSD test (α = 0.05). Pairwise genetic distances were calculated using the K2P model in MEGA 12.

## RESULTS

### Molecular Identification of Aphid Species Using COI Sequences

A total of eleven aphid DNA sequences were successfully analyzed using the NCBI BLAST tool against the GenBank database for species confirmation (Table 2).

**Table 2.**
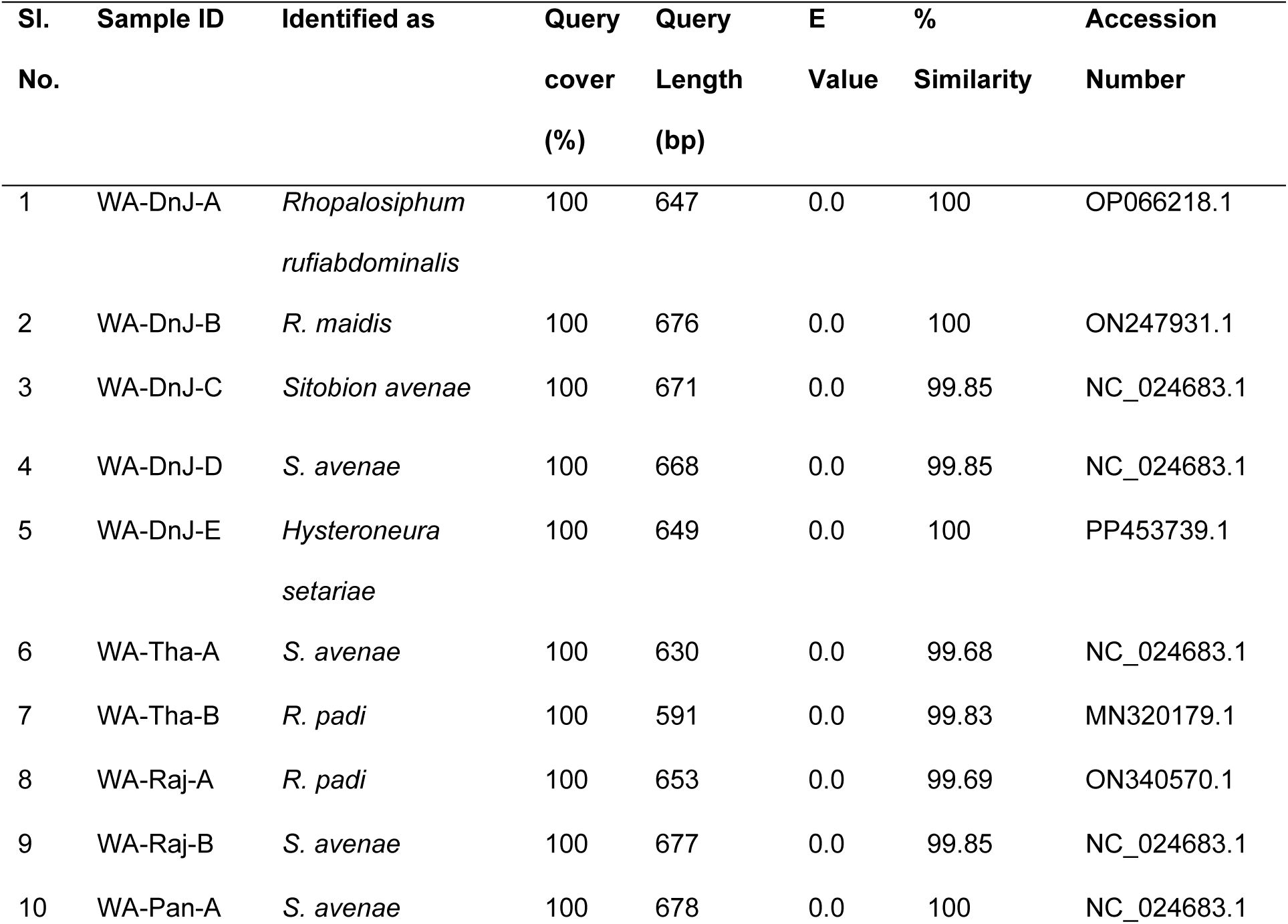
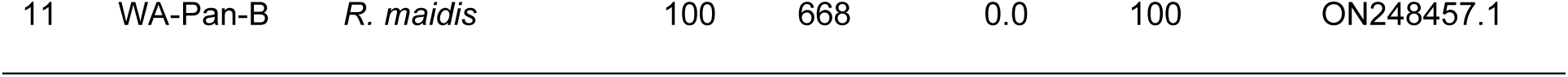
The percent similarity of the analyzed Aphid samples with GenBank accession number

**Table 3.** Pairwise evolutionary genetic distances among wheat aphid COI sequences based on the Kimura 2-parameter (K2P) model

| Sample | WA-DnJ-A | WA-DnJ-B | WA-DnJ-C | WA-DnJ-D | WA-DnJ-E | WA-Tha-A | WA-Tha-B | WA-Raj-A | WA-Raj-B | WA-Pan-A | WA-Pan-B |
| --- | --- | --- | --- | --- | --- | --- | --- | --- | --- | --- | --- |
| WA-DnJ-A | - |  |  |  |  |  |  |  |  |  |  |
| WA-DnJ-B | 0.0679 | - |  |  |  |  |  |  |  |  |  |
| WA-DnJ-C | 0.0969 | 0.0893 | - |  |  |  |  |  |  |  |  |
| WA-DnJ-D | 0.0969 | 0.0893 | 0.0000 | - |  |  |  |  |  |  |  |
| WA-DnJ-E | 0.0853 | 0.1034 | 0.1194 | 0.1194 | - |  |  |  |  |  |  |
| WA-Tha-A | 0.0969 | 0.0893 | 0.0000 | 0.0000 | 0.1194 | - |  |  |  |  |  |
| WA-Tha-B | 0.0491 | 0.0738 | 0.0891 | 0.0891 | 0.0814 | 0.0891 | - |  |  |  |  |
| WA-Raj-A | 0.0491 | 0.0738 | 0.0891 | 0.0891 | 0.0814 | 0.0891 | 0.0000 | - |  |  |  |
| WA-Raj-B | 0.0969 | 0.0893 | 0.0000 | 0.0000 | 0.1194 | 0.0000 | 0.0891 | 0.0891 | - |  |  |
| WA-Pan-A | 0.0969 | 0.0893 | 0.0000 | 0.0000 | 0.1194 | 0.0000 | 0.0891 | 0.0891 | 0.0000 | - |  |
| WA-Pan-B | 0.0660 | 0.0018 | 0.0914 | 0.0914 | 0.1054 | 0.0914 | 0.0758 | 0.0758 | 0.0914 | 0.0914 | - |

The BLAST results revealed the presence of five aphid species infesting wheat across the sampled locations: *Sitobion avenae*, *Rhopalosiphum padi*, *Rhopalosiphum maidis*, *Rhopalosiphum rufiabdominalis*, and *Hysteroneura setariae*. Among the identified species, *S. avenae* was the most prevalent, detected in five samples collected from Dinajpur, Thakurgaon, Rajshahi, and Panchagarh. Sequence similarity for *S. avenae* ranged from 99.68% to 100%, with query lengths varying between 630 and 678 bp and all sequences showing complete query coverage (100%) and an E-value of 0.0.

*Rhopalosiphum padi* was identified in two samples from Thakurgaon and Rajshahi, exhibiting 99.69–99.83% sequence similarity with query lengths ranging from 591 to 653 bp. Similarly, *R. maidis* was identified in two samples from Dinajpur and Panchagarh, both showing 100% similarity with query lengths of 668–676 bp. A single sample from Dinajpur was identified as *R. rufiabdominalis*, showing 100% similarity and a query length of 647 bp. Another Dinajpur sample was identified as *H. setariae* with 100% sequence similarity and a query length of 649 bp.

All sequences exhibited 100% query coverage and an E-value of 0.0, confirming strong alignment confidence and reliable species-level identification through COI barcode analysis. The obtained sequences matched reference accessions deposited in GenBank, are presented in Table 2.

### Phylogenetic Analysis of Aphid Species Based on COI Sequences

Phylogenetic analysis of the mitochondrial cytochrome c oxidase subunit I (COI) gene sequences confirmed the molecular identity of the aphid specimens and revealed clear clustering according to species (Fig 1).

**Fig 1.**
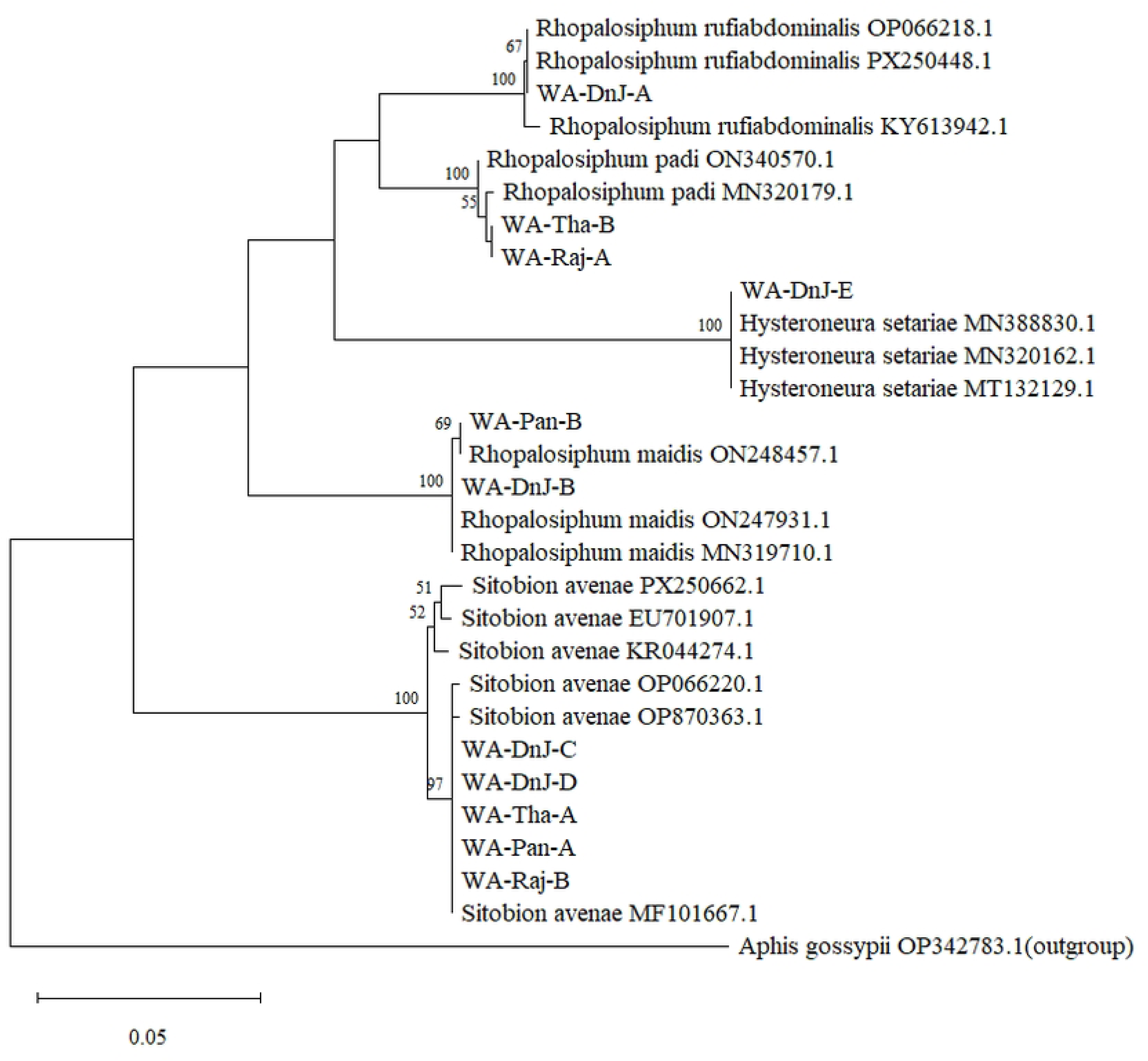
Phylogenetic tree inferred using the Maximum Likelihood (ML) method of eleven mtDNA CO1 region of wheat aphid sequences obtained from different locations of Bangladesh together with reference sequences. The tree is based on the Tamura-Nei model. The tree was resampled with 1000 bootstrap replicates. Bootstrap support values on the branches are given.

The Neighbor-Joining tree grouped the eleven obtained sequences into five distinct clades corresponding to *Sitobion avenae*, *Rhopalosiphum padi*, *Rhopalosiphum maidis*, *Rhopalosiphum rufiabdominalis*, and *Hysteroneura setariae*, in agreement with the BLAST identification results (Fig 1).

The *S. avenae* clade contained five sequences (WA-DnJ-C, WA-DnJ-D, WA-Tha-A, WA-Pan-A, and WA-Raj-B), which clustered closely with reference sequences from GenBank (PX250662.1, EU701907.1, KR044274.1, OP066220.1, OP870363.1, and MF101667.1). This clade was strongly supported with a bootstrap value of 97, indicating high genetic similarity among the isolates despite their geographic origin. The *R. padi* clade comprised two sequences (WA-Tha-B and WA-Raj-A), clustering with reference accessions ON340570.1 and MN320179.1, supported by moderate bootstrap values (55–100). The *R. maidis* sequences (WA-DnJ-B and WA-Pan-B) grouped with GenBank reference sequences ON247931.1, ON248457.1, and MN319710.1, showing strong bootstrap support (100), although slight divergence was observed within the clade.

A single isolate, WA-DnJ-A, formed a distinct cluster with *R. rufiabdominalis* reference sequences (OP066218.1, PX250448.1, and KY613942.1), supported by a bootstrap value of 100, confirming its taxonomic identity. Likewise, WA-DnJ-E clustered with *H. setariae* reference sequences (MN388830.1, MN320162.1, and MT132129.1) with strong bootstrap support (100).

The phylogenetic tree clearly separated all identified aphid species into species-specific lineages, while the outgroup species, *Aphis gossypii* (OP342783.1), formed a distant branch, validating the robustness of the phylogenetic reconstruction (Fig 1).

### Evolutionary divergence analysis

Pairwise genetic distance analysis based on the Kimura 2-parameter model revealed clear differentiation among wheat aphid species. Intraspecific divergence was very low, ranging from 0.0 to 0.18%, whereas interspecific divergence ranged from 4.9 to 11.9%. The highest genetic divergence was observed between *Sitobion avenae* and *Hysteroneura setariae* (11.9%), while the lowest interspecific divergence was recorded between *Rhopalosiphum padi* and *Rhopalosiphum rufiabdominalis* (4.9%). Within species, no variation was observed among sequences of *Sitobion avenae* and *Rhopalosiphum padi*, indicating identical haplotypes across different geographic locations. A low level of variation (0.18%) was detected within *Rhopalosiphum maidis*.

### Morphological comparison of aphid species infesting wheat

#### General features of live specimens

The morphology of *Sitobion avenae* was characterized by a broadly spindle-shaped body varying in color from yellowish-green to reddish-brown, accompanied by faint intersegmental markings. The species possessed six-segmented black antennae and elongated dark cornicles, which served as important diagnostic features (Fig 2).

**Fig 2.**
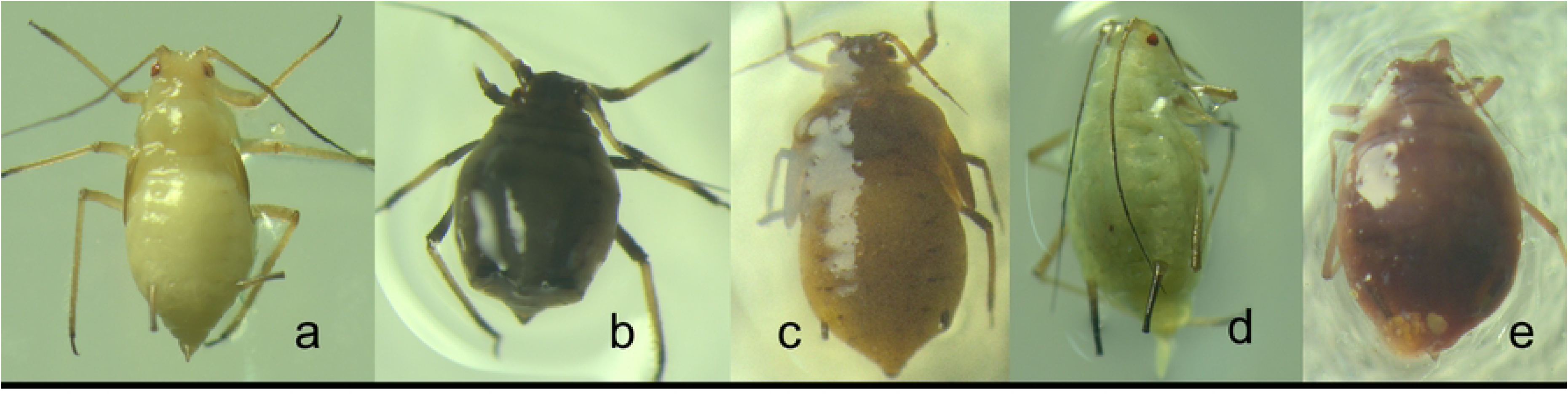
General morphological features of live specimens of the five wheat aphid species identified in wheat fields of Bangladesh: (a) *Sitobion avenae*; (b) *Hysteroneura setariae*; (c) *Rhopalosiphum maidis*; (d) *Rhopalosiphum padi*; and (e) *Rhopalosiphum rufiabdominalis*.

*Hysteroneura setariae* was distinguished by a broadly oval body with a dark chocolate-brown coloration and a slight olive tint. Morphologically, the species exhibited relatively shorter appendages and a compact body form (Fig 2). Specimens identified as *Rhopalosiphum maidis* were small and elongated, exhibiting a dark olive-green body coloration. A prominent diagnostic characteristic was the darker pigmented region surrounding the base of the cornicles. Morphometric examination revealed six-segmented antennae, slightly swollen black cornicles, a short dark cauda, and tubercles with a straight unguis (Fig 2).

*Rhopalosiphum padi* was characterized by a broadly oval olive-green body with distinctive reddish pigmentation at the bases of the cornicles. Both the cornicles and cauda were darkly pigmented, providing useful distinguishing features for species identification (Fig 2).

The specimens identified as *Rhopalosiphum rufiabdominalis* exhibited a broadly oval body shape with olive-green coloration and distinct reddish pigmentation surrounding the cornicle bases. Diagnostic morphological features included five-segmented antennae, dark cornicles, and a black cauda that was conspicuously shorter than the cornicles (Fig 2).

#### Key morphological traits of mounted specimens

Significant interspecific variation was observed among the five aphid species for all measured morphological characteristics, including body length, antenna length, tubercle width, cauda length, and cornicle length (ANOVA; *P* < 0.0001 for all traits) (Table 4).

**Table 4.**
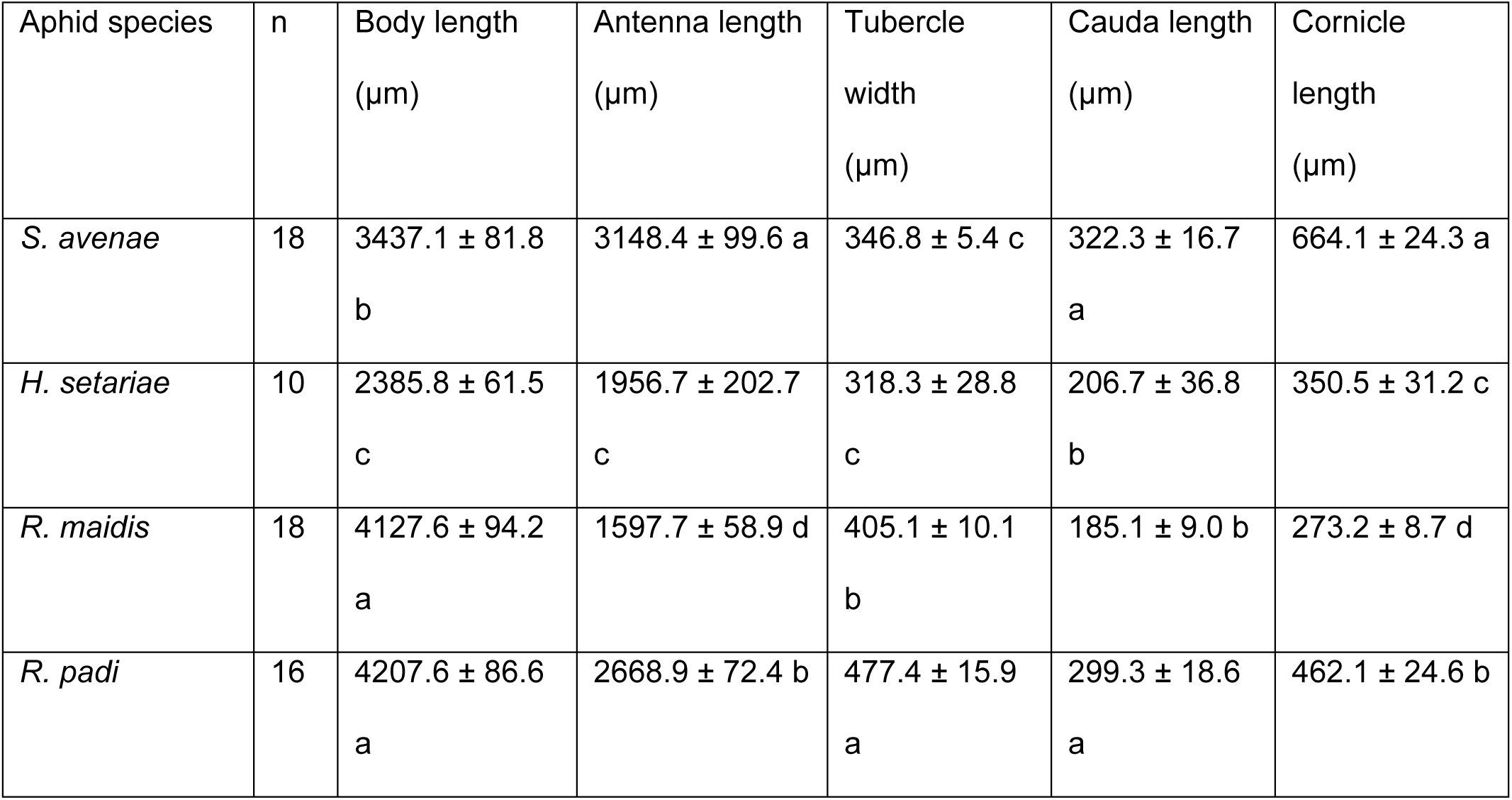
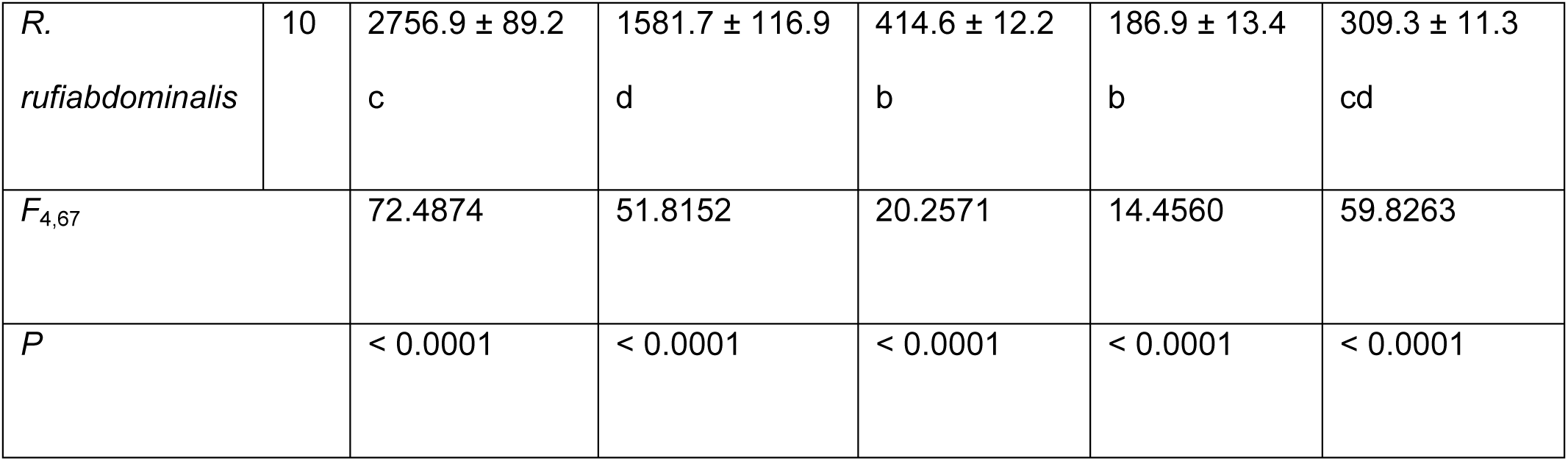
Key morphological traits (mean ± SE) of aphid species infesting wheat in Bangladesh

| Aphid species | n | Body length<br>($\mu\text{m}$) | Antenna length<br>($\mu\text{m}$) | Tubercle<br>width<br>($\mu\text{m}$) | Cauda length<br>($\mu\text{m}$) | Cornicle<br>length<br>($\mu\text{m}$) |
| --- | --- | --- | --- | --- | --- | --- |
| *S. avenae* | 18 | 3437.1 ± 81.8<br>b | 3148.4 ± 99.6 a | 346.8 ± 5.4 c | 322.3 ± 16.7<br>a | 664.1 ± 24.3 a |
| *H. setariae* | 10 | 2385.8 ± 61.5<br>c | 1956.7 ± 202.7<br>c | 318.3 ± 28.8<br>c | 206.7 ± 36.8<br>b | 350.5 ± 31.2 c |
| *R. maidis* | 18 | 4127.6 ± 94.2<br>a | 1597.7 ± 58.9 d | 405.1 ± 10.1<br>b | 185.1 ± 9.0 b | 273.2 ± 8.7 d |
| *R. padi* | 16 | 4207.6 ± 86.6<br>a | 2668.9 ± 72.4 b | 477.4 ± 15.9<br>a | 299.3 ± 18.6<br>a | 462.1 ± 24.6 b |
| *R. rufiabdominalis* | 10 | 2756.9 ± 89.2<br>c | 1581.7 ± 116.9<br>d | 414.6 ± 12.2<br>b | 186.9 ± 13.4<br>b | 309.3 ± 11.3<br>cd |
| $F_{4,67}$ | | 72.4874 | 51.8152 | 20.2571 | 14.4560 | 59.8263 |
| *P* |  | < 0.0001 | < 0.0001 | < 0.0001 | < 0.0001 | < 0.0001 |

The body lengths of mounted specimens are shown in figure 3, where *R. padi* (4207.6 ± 86.6 μm) exhibited the greatest body length, which was statistically similar to *R. maidis* (4127.6 ± 94.2 μm). In contrast, *H. setariae* showed the shortest body length (2385.8 ± 61.5 μm), which was statistically comparable to *R. rufiabdominalis* (2756.9 ± 89.2 μm). *S. avenae* (3437.1 ± 81.8 μm) exhibited an intermediate body size between the larger and smaller species groups (Table 4).

**Fig 3.**
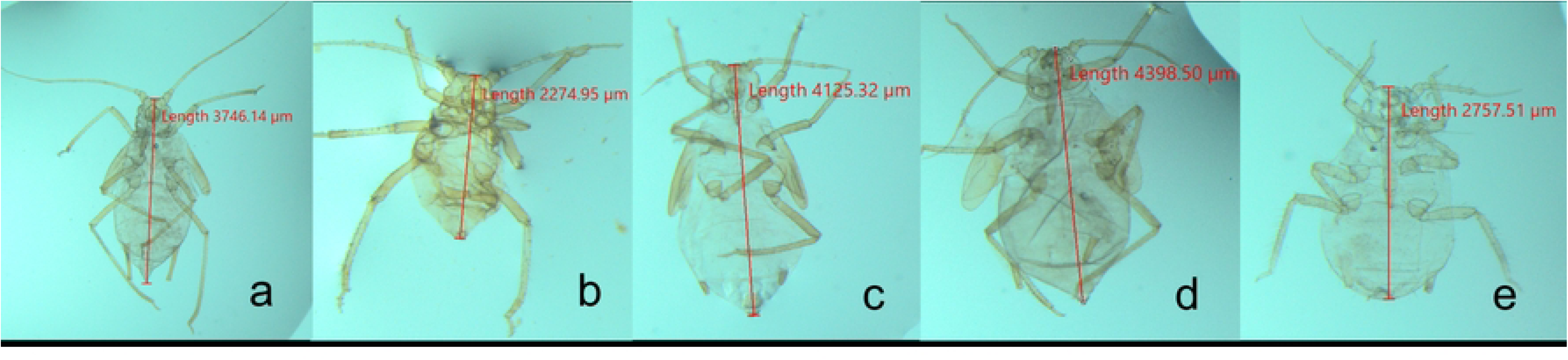
Body length of mounted specimens of the five wheat aphid species identified in wheat fields of Bangladesh: (a) *Sitobion avenae*; (b) *Hysteroneura setariae*; (c) *Rhopalosiphum maidis*; (d) *Rhopalosiphum padi*; and (e) *Rhopalosiphum rufiabdominalis*.

Antenna lengths of mounted specimens are shown in figure 4, where *S. avenae* recorded the longest antennae (3148.4 ± 99.6 μm), followed by *R. padi* (2668.9 ± 72.4 μm). In contrast, *R. maidis* (1597.7 ± 58.9 μm) and *R. rufiabdominalis* (1581.7 ± 116.9 μm) possessed the shortest antennae and were statistically comparable, whereas H. setariae showed intermediate values (1956.7 ± 202.7 μm) (Table 4).

**Fig 4.**
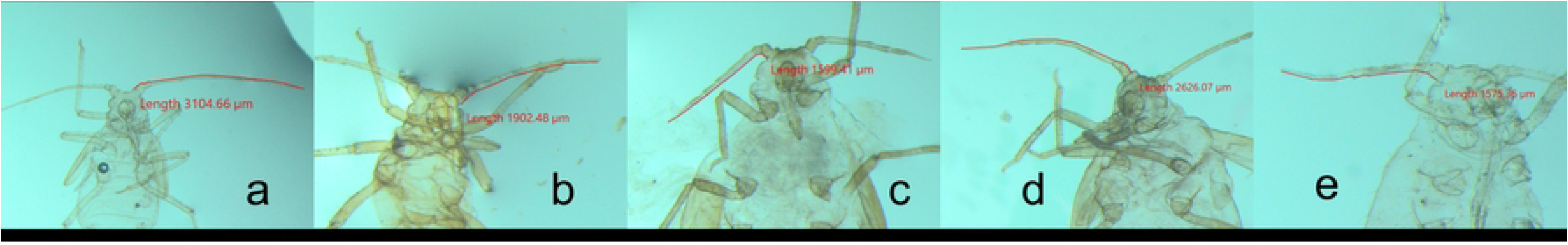
Antenna length of mounted specimens of the five wheat aphid species identified in wheat fields of Bangladesh: (a) *Sitobion avenae*; (b) *Hysteroneura setariae*; (c) *Rhopalosiphum maidis*; (d) *Rhopalosiphum padi*; and (e) *Rhopalosiphum rufiabdominalis*.

Tubercle widths are shown in figure 5, where *R. padi* exhibited the widest tubercles (477.4 ± 15.9 μm), followed by *R. rufiabdominalis* (414.6 ± 12.2 μm) and *R. maidis* (405.1 ± 10.1 μm), which were statistically similar. The narrowest tubercles were observed in *S. avenae* (346.8 ± 5.4 μm) and *H. setariae* (318.3 ± 28.8 μm) (Table 4).

**Fig 5.**
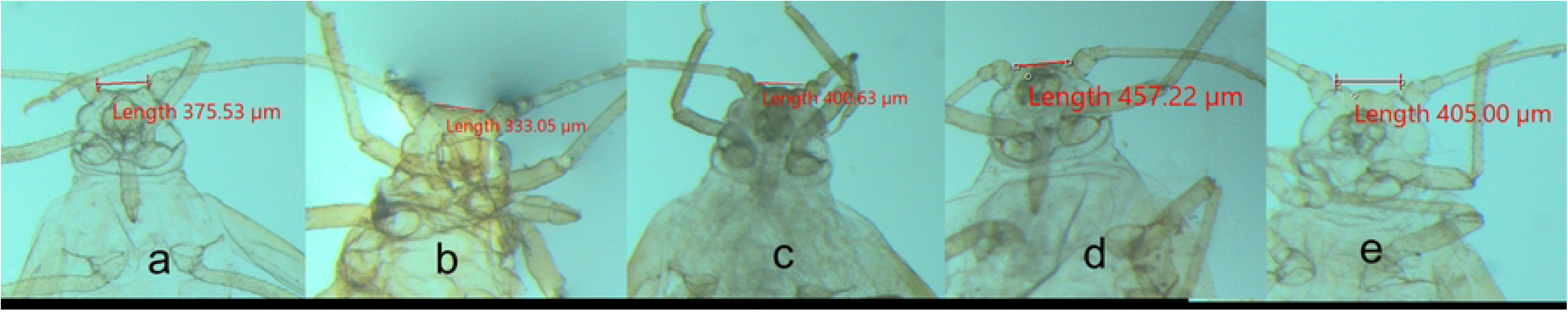
Tubercle width of mounted specimens of the five wheat aphid species identified in wheat fields of Bangladesh: (a) *Sitobion avenae*; (b) *Hysteroneura setariae*; (c) *Rhopalosiphum maidis*; (d) *Rhopalosiphum padi*; and (e) *Rhopalosiphum rufiabdominalis*.

Caudae lengths are displayed in figure 6, where *S. avenae* (322.3 ± 16.7 μm) and *R. padi* (299.3 ± 18.6 μm) had significantly longer caudae than the other species, while *R. maidis* (185.1 ± 9.0 μm), *R. rufiabdominalis* (186.9 ± 13.4 μm), and *H. setariae* (206.7 ± 36.8 μm) showed comparatively shorter cauda lengths (Table 4).

**Fig 6.**
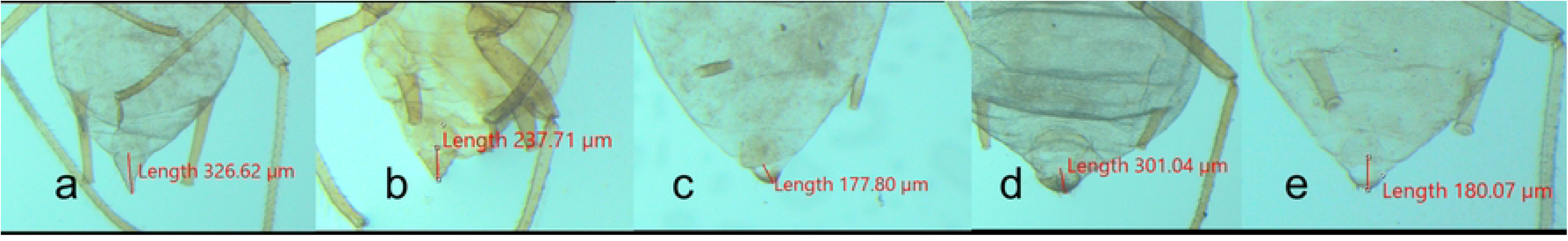
Cauda length of mounted specimens of the five wheat aphid species identified in wheat fields of Bangladesh: (a) *Sitobion avenae*; (b) *Hysteroneura setariae*; (c) *Rhopalosiphum maidis*; (d) *Rhopalosiphum padi*; and (e) *Rhopalosiphum rufiabdominalis*.

Cornicle lengths are demonstrated in figure 7, where *S. avenae* possessed significantly longer cornicles (664.1 ± 24.3 μm) than all other species, followed by *R. padi* (462.1 ± 24.6 μm). The shortest cornicles were recorded in *R. maidis* (273.2 ± 8.7 μm), while *R. rufiabdominalis* (309.3 ± 11.3 μm) and *H. setariae* (350.5 ± 31.2 μm) exhibited intermediate values (Table 4).

**Fig 7.**
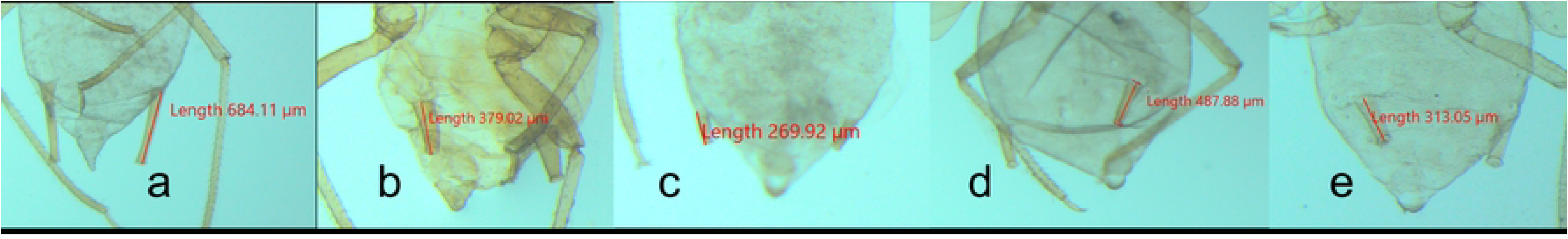
Cornicle length of mounted specimens of the five wheat aphid species identified in wheat fields of Bangladesh: (a) *Sitobion avenae*; (b) *Hysteroneura setariae*; (c) *Rhopalosiphum maidis*; (d) *Rhopalosiphum padi*; and (e) *Rhopalosiphum rufiabdominalis*.

## DISCUSSION

The present study provides the first comprehensive integrative assessment of wheat aphid diversity in Bangladesh by combining mitochondrial COI DNA barcoding with detailed morphological characterization. This approach enabled the reliable identification of five aphid species associated with wheat in Bangladesh, namely *Sitobion avenae*, *Rhopalosiphum padi*, *R. maidis*, *R. rufiabdominalis*, and *Hysteroneura setariae*. Although these species have been reported from neighboring countries and other wheat-producing regions worldwide [8,13,29,30], information regarding their occurrence and distribution in Bangladesh has remained limited and largely based on morphological observations. Our findings therefore establish an updated species inventory and provide the first molecular confirmation of wheat-infesting aphids in Bangladesh, forming an essential baseline for future ecological, epidemiological, and pest management studies.

DNA barcoding using the mitochondrial COI gene proved highly effective for species discrimination [17,31,32], and in this study all sequences exhibiting 99.68–100% similarity to reference sequences in GenBank and complete query coverage. The clear correspondence between BLAST results and phylogenetic clustering demonstrates the reliability of COI as a molecular marker for species identification. Similar studies have shown that COI barcoding provides accurate species delimitation for agriculturally important aphids [28,33,34], particularly where morphological variation or phenotypic plasticity complicates conventional taxonomy [35,36,37,38]. The present results further confirm that DNA barcoding is a valuable complementary tool for routine identification of cereal aphids, especially during immature stages or when specimens are damaged during collection.

Phylogenetic analysis further validated the taxonomic assignments by grouping all Bangladeshi specimens into well-supported species-specific clades together with their corresponding reference sequences. The high bootstrap support observed for most clades indicates strong genetic consistency within species despite collections from geographically separated wheat-growing regions. Moreover, the absence of distinct geographic clustering among isolates of the same species suggests substantial gene flow or recent dispersal across growing areas [39,40]. Such genetic homogeneity has also been reported in aphid populations, where long-distance migration aided by prevailing winds and repeated colonization events contributes to low regional genetic differentiation [41,42,43,44]. The relatively limited intraspecific variation detected in this study therefore appears consistent with the high dispersal capacity and predominantly parthenogenetic reproduction that characterize many cereal aphids [45,46,47,48].

Pairwise genetic distance analysis further demonstrated the presence of a clear DNA barcode gap, with intraspecific divergence remaining below 0.2% while interspecific divergence ranged from approximately 5% to 12%. This distinct separation supports the robustness of COI for discriminating closely related aphid taxa and agrees with previous DNA barcoding studies reporting substantially higher genetic divergence among species than within species [49,50,51]. The complete absence of sequence variation among several *S. avenae* and *R. padi* isolates indicates the occurrence of identical haplotypes across multiple wheat-growing regions, whereas the slight divergence observed within *R. maidis* may reflect localized genetic variation or the presence of multiple maternal lineages. Although the current sampling size was limited, these findings suggest that cereal aphid populations in Bangladesh possess relatively low mitochondrial diversity.

Among the identified species, *S. avenae* was the most frequently detected aphid, occurring across multiple wheat-growing districts. This widespread distribution is consistent with reports from Europe, China, India, and other wheat-producing regions where *S. avenae* is regarded as one of the dominant cereal aphids [52,53,54,55]. Its prevalence in Bangladesh may reflect its broad ecological adaptability, efficient reproductive capacity, and strong preference for wheat as a host plant [56,57,58]. In contrast, *R. padi* and *R. maidis* were detected less frequently but occurred across multiple locations, suggesting that both species contribute to the wheat aphid complex rather than acting as occasional incidental pests [59,60]. The detection of *R. rufiabdominalis* and *H. setariae* only in Dinajpur indicates that these species may possess more restricted distributions, occur at lower population densities, or utilize alternative host plants before colonizing wheat [61,62].

Morphological examinations corroborated the molecular identifications and demonstrated that several morphometric characters possess strong taxonomic value for separating cereal aphid species. Significant interspecific differences in body length, antenna length, cornicle length, cauda length, and tubercle width indicate that these characters remain reliable diagnostic traits when specimens are properly mounted and examined [63,64]. Particularly informative were the exceptionally long cornicles and antennae of *S. avenae*, the relatively large body size of *R. padi* and *R. maidis*, and the shorter appendages observed in *H. setariae* and *R. rufiabdominalis* [13,65,66]. These findings are largely consistent with classical taxonomic descriptions and regional identification keys developed for cereal aphids. The strong agreement between molecular and morphological datasets demonstrates the advantages of integrative taxonomy, reducing the likelihood of misidentification that may arise when either approach is used independently.

Accurate identification of wheat aphid species has important implications for integrated pest management in Bangladesh. Different aphid species vary in host preference, seasonal occurrence, reproductive biology, insecticide susceptibility, and efficiency as vectors of economically important barley and cereal yellow dwarf viruses [5,38,67–70]. Consequently, effective pest management requires reliable species identification before monitoring programs, economic threshold development, or biological control strategies can be optimized. The molecular reference sequences generated in this study provide valuable resources for future diagnostic programs, while the accompanying morphological descriptions facilitate rapid field identification by researchers and extension personnel who may not have access to molecular facilities.

Although this study substantially advances current knowledge of wheat aphid diversity in Bangladesh, several limitations should be acknowledged. Sampling was conducted from a limited number of wheat-producing regions and included relatively few specimens for each species. Consequently, the present study likely represents only part of the country’s aphid diversity. In addition, only the mitochondrial COI marker was analyzed, which reflects maternal inheritance and may not fully capture population-level genetic variation. Future studies incorporating larger sample sizes, additional ecological zones, seasonal sampling, and high-resolution molecular markers such as microsatellites, SNPs, or genome-wide sequencing would provide deeper insights into aphid population genetics, migration dynamics, host adaptation, and insecticide resistance. Furthermore, investigations linking aphid species composition with virus transmission efficiency, natural enemy communities, and climatic variables would contribute substantially to the development of sustainable wheat pest management strategies in Bangladesh.

In conclusion, this study presents the first integrative molecular and morphological framework for the identification of five wheat aphid species in Bangladesh: *Sitobion avenae*, *Rhopalosiphum padi*, *R. maidis*, *R. rufiabdominalis*, and *Hysteroneura setariae*. The strong concordance between DNA barcoding, phylogenetic analyses, and morphological characterization demonstrates the robustness and reliability of the integrative taxonomic approach adopted in this study. These findings establish a solid taxonomic baseline for future investigations of wheat aphid diversity, population genetics, and invasion dynamics. Moreover, the baseline data generated here will support enhanced biodiversity assessments, pest surveillance, insecticide resistance monitoring, and the development of evidence-based integrated pest management (IPM) strategies to safeguard wheat production in Bangladesh and across South-East Asia under changing agricultural and climatic conditions.

## ACKNOWLEDGEMENTS

The authors gratefully acknowledge the Ministry of Agriculture, Government of the People’s Republic of Bangladesh, for its continued support of agricultural research. This work was made possible through the establishment and strengthening of laboratory facilities under several collaborative projects and funding initiatives, including the USAID Bangladesh Fighting Fall Armyworm Project and the Feed the Future Integrated Pest Management Activity, both implemented by CIMMYT. Additional support was provided through the AFACI Asia Regional FAW and BPH Diagnostics, Monitoring, and Surveillance Program. The authors sincerely appreciate the technical and institutional support provided by these organizations, which contributed significantly to the successful completion of this study.

